# Is microbial activity in tropical forests limited by phosphorus availability? Evidence from a tropical forest in China

**DOI:** 10.1101/575001

**Authors:** Taiki Mori, Xiankai Lu, Cong Wang, Qinggong Mao, Senhao Wang, Wei Zhang, Jiangming Mo

## Abstract

The prevailing paradigm for soil microbial activity in tropical forests is that microbial activity is limited by phosphorus (P) availability. Thus, exogenous P addition should increase rates of organic matter decomposition. Studies have also confirmed that soil respiration is accelerated when P is added experimentally. However, we hypothesize that the increased rates of soil microbial respiration could be due to the release of organic material from the surface of soil minerals when P is added, because P is more successful at binding to soil particles than organic compounds. In this study, we demonstrate that P addition to soil is associated with significantly higher dissolved organic carbon (DOC) content in a tropical evergreen forest in southern China. Our results indicate that P fertilization stimulated soil respiration but suppressed litter decomposition. Results from a second sorption experiment revealed that the recovery ratio of added DOC in the soil of a plot fertilized with P for 9 years was larger than the ratio in the soil of a non-fertilized plot, although the difference was small. We also conducted a literature review on the effects of P fertilization on the decomposition rates of litter and soil organic matter at our study site. Previous studies have consistently reported that P addition led to higher response ratios of soil microbial respiration than litter decomposition. Therefore, experiments based on P addition cannot be used to test whether microbial activity is P-limited in tropical forest soils, because organic carbon desorption occurs when P is added. Our findings suggest that the prevailing paradigm on the relationship between P and microbial activity in tropical forest soils should be re-evaluated.

## Introduction

In recent decades, nutrient inputs into ecosystems have been greatly altered by human activities (Gallardo and Schlesinger, 1994; Galloway et al., 2004). A complete understanding of how nutrient limitations impact soil microbial activity, which plays an important role in the decomposition of organic carbon (C), is essential for predicting global C cycling. Traditionally, soil microbial activity in tropical forests is considered to be limited by phosphorus (P) availability. Thus, the addition of exogenous P is thought to stimulate organic matter decomposition. Accordingly, a number of studies have demonstrated that the respiration rates of soil microbes increase with the experimental addition of P (Cleveland et al., 2002; Duah-Yentumi et al., 1998; Gnankambary et al., 2008; Ilstedt and Singh, 2005; Mori et al., 2016). Hence, these studies concluded that microbial activity in tropical forest soils is P-limited.

However, Mori et al. (2018) suggest that the increased soil microbial respiration may be due to the release of organic matter from soil mineral surfaces, because P competes more successfully for binding sites on soil particles than many organic compounds when added artificially (Kaiser & Zech 1996; Afif et al. 1995; McKercher, and Anderson, 1989; Celi and Barberis, 2005; Guppy et al., 2005; Ruttenberg and Sulak, 2011). The release of organic matter would increase the amount of resources available to soil microbes and stimulate microbial respiration. Thus, soil microbial activity is predominantly C-limited, not P-limited. As such, the P-fertilization method, which is commonly used to test microbial P-limitation, would overestimate the impacts of P addition on the decomposition of soil organic matter. One would then conclude that P addition would increase the decomposition rates of soil organic matter (with mineral soil) more than the decomposition rates of litter (without mineral soil). As expected, most of the studies that reported stimulatory effects of P addition on microbial respiration investigated soils in tropical ecosystems (Cleveland et al., 2002; Galicia and Garcia-Oliva, 2004; Mori et al., 2013b; Mori et al., 2016), whereas studies on litter decomposition reported that P addition had neutral (Barantal et al., 2012; Cleveland et al., 2006; Davidson et al., 2004; McGroddy et al., 2004) or negative effects (Chen et al., 2013; Mori et al., 2015; Zheng et al., 2017).

Our earlier work also revealed that experimental P addition stimulated soil respiration in the field (Liu et al., 2012) and suppressed leaf litter decomposition (Chen et al., 2013; Fig. 1). As discussed above, the difference in impacts may be due to the increase in C availability following experimental P addition. In this study, we investigated whether P addition desorbs dissolved organic C (DOC) from solid soil particles in the same tropical forest where we conducted previous work (Chen et al., 2013; Liu et al., 2012; Fig. 1). We also conducted a literature review on the impacts of P fertilization on the decomposition of soil organic matter and litter. We hypothesized that the response ratio of soil microbial respiration to experimental P addition would be larger than that of litter decomposition.

**Figure 1.**
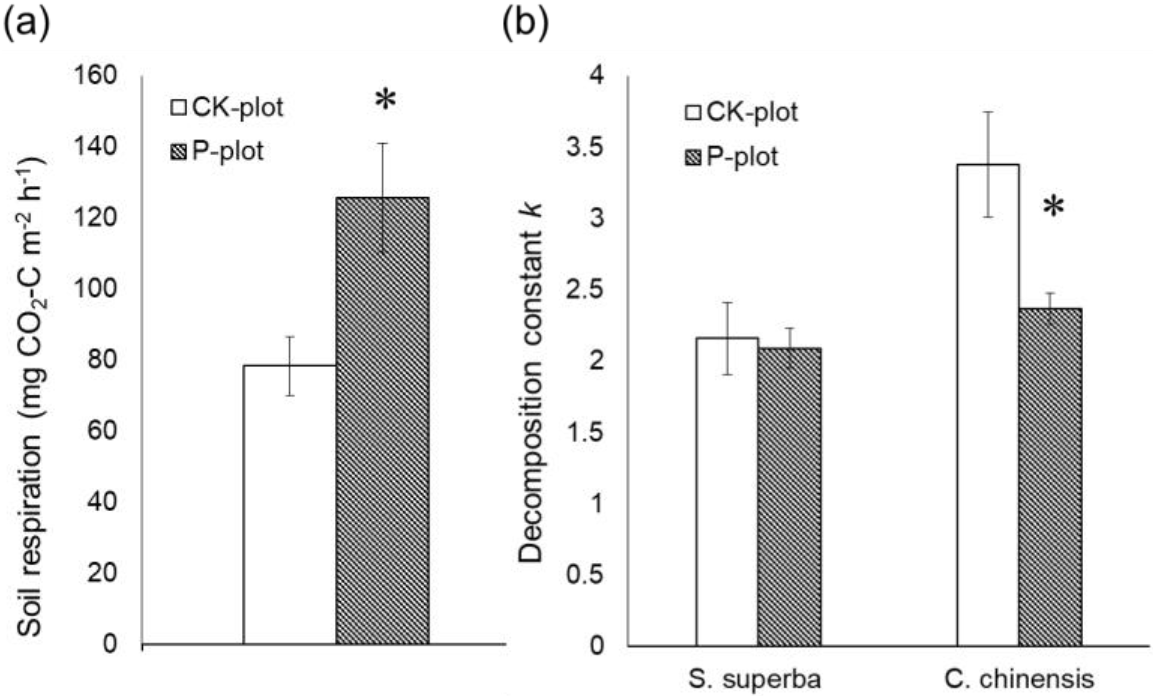
Effects of phosphorus (P) fertilization in the field on (a) soil respiration (Liu et al., 2012) and (b) decomposition of litter from *Schima superba* Chardn. & Champ. and *Castanopsis chinensis* Hance (Chen et al., 2013). P fertilization increased soil respiration rates, but had negative or no effects on litter decomposition. Soil respiration rates were averaged over three months (July to September 2009). Decomposition rates (*k*) were determined by running a single negative exponential model on 18 months of decomposition data. CK-plot, control plot without P fertilization; P-plot, treatment plot fertilized with 150 kg P·ha^-1^·yr^-1^ since Feb. 2007. Error bars indicate standard errors (SE). * *P* < 0.05 (ANOVA). Additional details can be found in Liu et al. (2012) and Chen et al. (2013).

## Materials and methods

### Site description

Our study was carried out in an old-growth monsoonal primary forest dominated by evergreen broadleaf trees, located in the Dinghushan Biosphere Reserve in the middle of Guangdong Province, southern China (112°10’E, 23°10N). This region experiences a subtropical monsoon climate, and the soil consists of lateritic red earth (oxisols) formed from sandstone (Mo et al., 2003). The mean annual precipitation is 1,927 mm; 75% of the rainfall occurs from March to August and only 6% occurs from December to February (Huang and Fan, 1982). The mean annual temperature is 21.0°C; July is the warmest month (28.0°C), and January is the coldest month (12.6°C). Results from ^14^C analysis revealed that this forest has not been anthropogenically disturbed for 400 years (Shen et al., 1999). The dominant tree species in this forest are *Castanopsis chinensis, Schima superba, Cryptocarya chinensis* (Hance) Hemsl., *Cryptocarya concinna, Machilus chinensis* (Champ. Ex Benth.) Hemsl., and *Syzygium rehderianum* Merr. & Perry, whereas the understory is dominated by *Calamus rhabdicladus* Burret, *Ardisia quinquegona* Bl., and *Hemigramma decurrens* (Hook.) Copel (Mo et al., 2003; Zhou et al., 2018). The characteristics of the study site are summarized in Table 1.

**Table 1.**
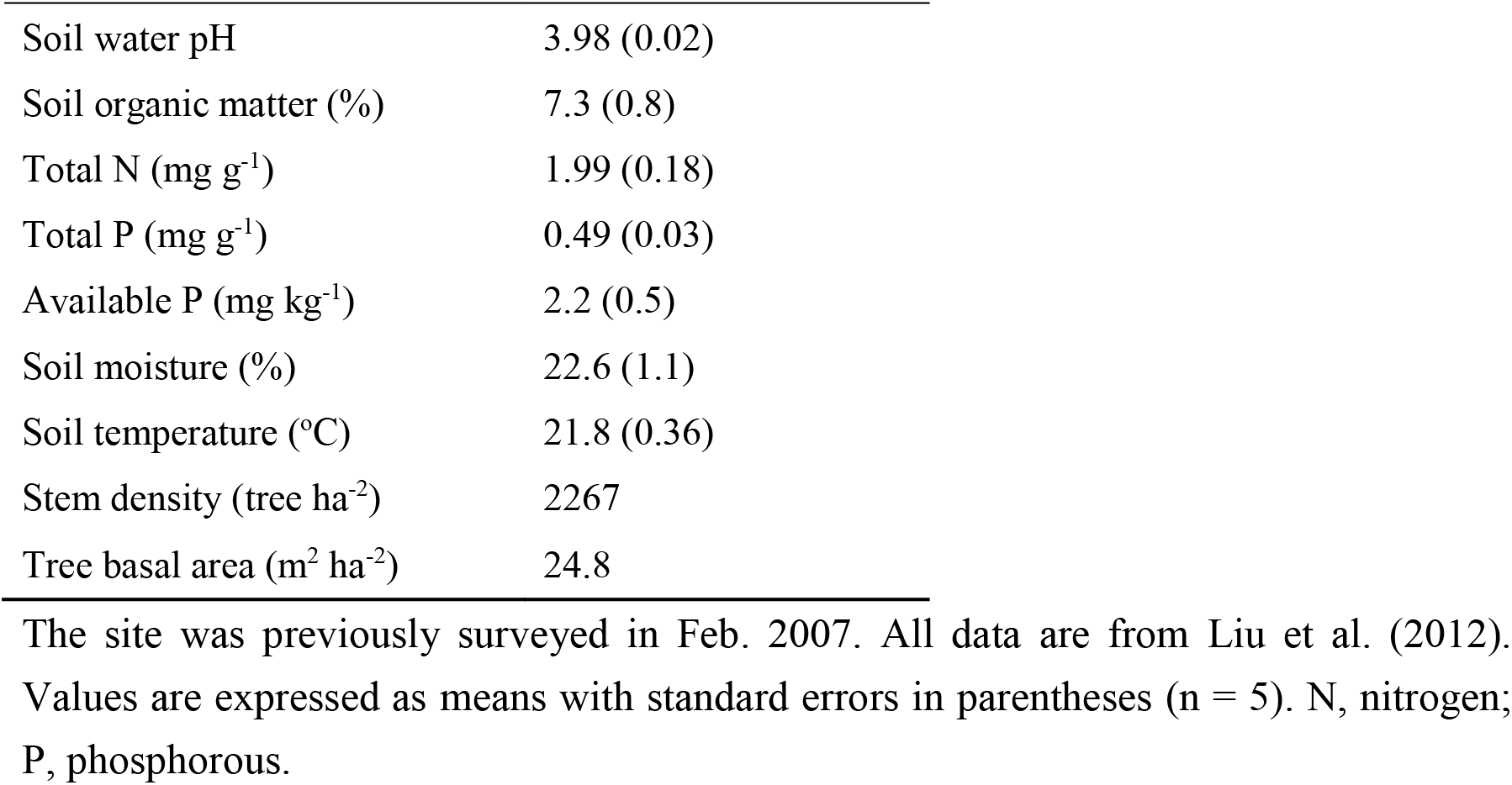
Characteristics of the study site.

### Long-term fertilization experiment

A long-term P fertilization experiment was initiated in 2007 (Liu et al., 2012), and five treatment (P-plot) and five control (CK-plot) plots were established. Each plot had an area of 5 × 5 m and was separated from other plots by buffer strips. Monosodium phosphate (NaH_2_PO_4_) solution (15 g P·m^-2^·yr^-1^) was used to add P to the treatment plots. The P fertilizer was mixed with 5 L of water and sprayed below the tree canopy using a backpack sprayer. CK-plots were sprayed with 5 L of water. Our experimental design, including plot size and fertilizer dosage, followed the protocol of a similar fertilization experiment in Costa Rica (Cleveland and Townsend, 2006).

### Sampling and laboratory experiment

Surface soil samples (depth = 10 cm) were collected in October 2016. We took six soil cores (3-cm inner diameter) at each plot and combined the six cores into a composite sample. The roots were removed from each soil sample, which was then passed through a 2-mm mesh sieve and mixed well. Leaf litter was collected from the unfertilized area to extract dissolved organic matter (DOM). We mixed 150 g of litter with 1,500 mL deionized water for 24 h, and extracted DOM by filtering the mixture through a 0.45-μm membrane filter (Merida, USA).

The soil collected from the CK-plots and P-plots were used in laboratory sorption experiments. We tested the effect of P addition on levels of water-extracted DOC in the soil from the CK-plots with and without additional DOM. First, soil samples were oven-dried. Then, 5 g of soil was shaken with monopotassium phosphate (KH2PO4; 0, 20, 100, or 500 μg P · g soil^-1^) and 3 mL DOM for 30 min under cool conditions. Thereafter, samples were stored in a refrigerator at 4°C to prevent microbial decomposition. Subsequently, samples were centrifuged, filtered through 0.45-μm membrane filters, and frozen.

We also tested the effects of long-term (9 years) P fertilization on the DOC recovery ratio (RR) in the field with externally added DOC. For this experiment, 3 mL of DOM was added to soils sampled from the CK-plots and P-plots, and DOC was extracted as described above. The DOC recovery ratio (%) was calculated as follows:

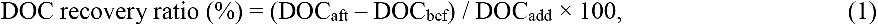

where DOC_aft_ is the concentration of DOC extracted from soil with added DOM, DOC_bef_ is the concentration of DOC extracted from soil without added DOM, and DOC_add_ is the concentration of DOC in the DOM added to the soil. Dissolved nitrogen (DN) content and recovery ratio were also determined in the same manner. DOC and DN content were determined using the Shimadzu total organic carbon analyzer (Shimadzu Corporation, Kyoto, Japan).

### Literature review

We searched for publications related to the effects of P fertilization on the decomposition rates of litter and soil organic matter at our study site using Web of Science. The following key words and combinations were used in the search: (“microbial respiration” OR (soil decomposition incubation) OR “litter bag” OR “organic matter decomposition” OR “litter decomposition”) AND (“phosph* add*” OR “P add*” OR “phosph* elevat*” OR “P elevat*” OR “phosph* fertiliz*” OR “P fertiliz*” OR “phosph* appl*” OR “P appl*”OR “phosph* enrich*” OR “P enrich*”).

We did not include studies conducted in streams, wetlands, or mangroves, or studies with field measurements of soil respiration data, as these data are affected by changes in root respiration rates and litter fall inputs from P addition. Additionally, studies on the effects of simultaneous additions of C and P on microbial respiration were excluded, because we were unable to separate out the P effects on the decomposition of soil organic matter (Nottingham et al., 2015). We also excluded data on soil microbial respiration from incubation experiments using soil taken from P-fertilized and unfertilized sites, because organic matter content could have been affected by fertilization in the field, and may have differed from the initial values estimated from the incubation samples. The means of data that were only reported in figures were extracted using DataThief (https://datathief.org/). For studies with multiple P doses or multiple sampling dates, we considered this repetition to be dependent (Mori, 2017), and only used the measurements corresponding to the highest doses or the latest sampling period. Multiple independent experiments within the same study, such as experiments conducted at different locations, were considered separately.

### Calculations and statistical analyses

The response ratio (RR) of soil microbial respiration or litter decomposition was calculated as follows:

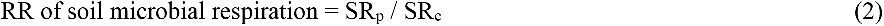

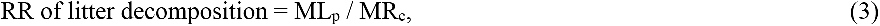

where SR_p_ is the rate of soil microbial respiration in soil with added P, SR_c_ is rate of soil microbial respiration in control soil, ML_p_ is litter mass loss (or carbon dioxide [CO_2_] emissions during decomposition) when P is added, and MR_c_ is litter mass loss when no P is added (control).

DOC and DN content and RRs were compared among treatment and control groups using one-way analysis of variance (ANOVA). R v.3.4.4 was used for all statistical analyses (R Core Team, 2018).

## Results and discussion

Our results revealed that the addition of P increased C and N availability. In the first sorption experiment, P addition to the soil in CK-plots significantly increased DOC and DN content (Fig. 2). In the second experiment, the recovery ratio of the added DOC was larger in the P-plots than in the CK-plots, but the difference was not significant (P = 0. 056, Fig. 3a). The recovery ratio of the added DN in the P-plots did not differ from the recovery ratio in the CK-plots (Fig. 3b). P addition likely causes the detachment of organic matter from soil mineral surfaces (Kaiser and Zech, 1996), which may have led to the higher DOC content and recovery ratios in this study. The DOC that was derived from the litter (e.g., leaves, branches, and roots) would be less adsorbed by the soil, and thus, more easily taken up by the microbes in the P-plots. This increased DOC uptake may have increased soil respiration rates (Liu et al. 2012). This mechanism would also explain why P fertilization stimulated soil respiration (Liu et al. 2012), but suppressed or had no effects on leaf litter decomposition (Chen et al. 2013; Fig. 1). In contrast, if the soil microbes were P-limited as the prevailing paradigm suggests, P fertilization would increase both soil respiration and litter decomposition rates. Inorganic P has been shown to detach organic compounds from mineral soils and accelerate soil microbial respiration even in P-rich soils (Spohn and Schleuss, in press). Similarly, the soil microbes in our study are likely to be C-limited rather than P-limited. Thus, the increase in the availability of abiotic C from P fertilization could have led to both an increase in soil respiration rates (Liu et al., 2012) and a decrease or lack of effect on litter decomposition rates (Chen et al., 2013). Nevertheless, the increased litter production and root respiration rates from P fertilization may have influenced our study outcomes as well.

**Figure 2.**
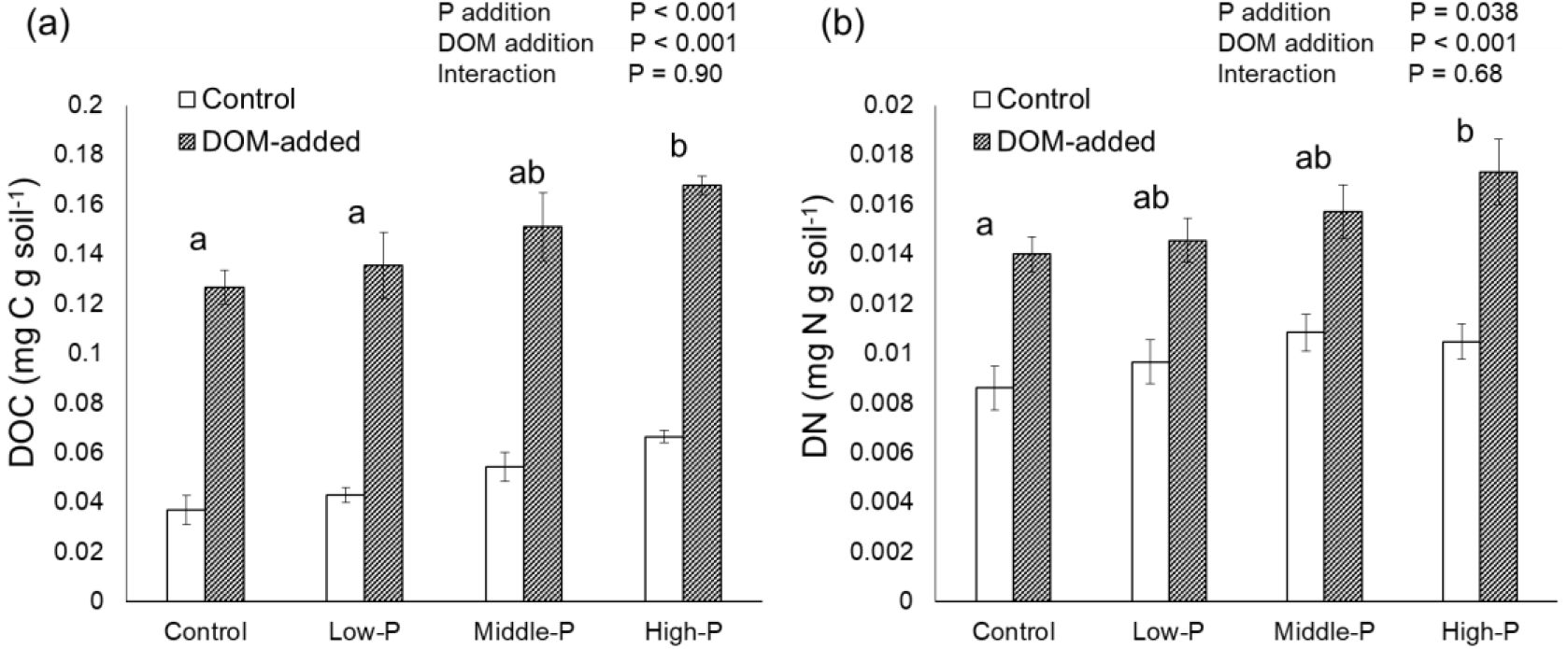
Effects of phosphorus (P) addition on water-extracted (a) dissolved organic carbon (DOC) and (b) dissolved nitrogen (DN). The P-values from one-way ANOVA are listed for each treatment. Different lowercase letters represent significant differences among P-fertilization treatments (*P* < 0.05, Tukey’s post-hoc test). Error bars indicate SE.

**Figure 3.**
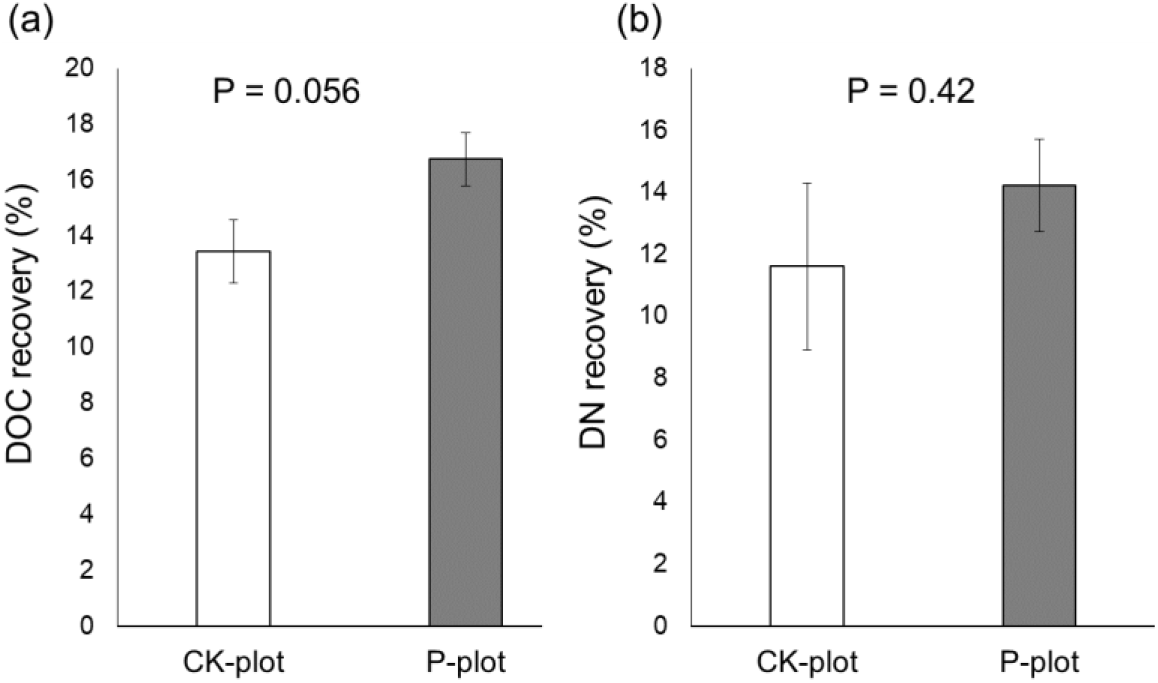
Effects of phosphorus (P) addition on the recovery of experimentally added (a) dissolved organic carbon (DOC) and (b) dissolved nitrogen (DN). P-values from one-way ANOVA are displayed in each plot. Error bars indicate SE.

From our literature review, other studies have also reported that P fertilization had contrasting effects on soil microbial respiration and litter decomposition. We did not include soil respiration data measured in the field, as litter input and root respiration could be affected by P fertilization. We analyzed the data from seven studies in total. The results from four experiments indicated that P addition resulted in increased rates of soil microbial respiration (Cleveland et al., 2002; Mori et al., 2010; Mori et al., 2016a). None of the seven studies reported that P addition increased litter decomposition rates (Cleveland et al., 2006; Hobbie and Vitousek, 2000; Mori et al., 2015). P addition in incubation experiments at a tropical tree plantation in south Sumatra, Indonesia significantly increased rates of soil microbial respiration (Mori et al., 2010), but decreased litter decomposition rates (Mori et al., 2015). Similarly, P fertilization in litter bag experiments had no effect on litter decomposition in a tropical primary rainforest in Costa Rica (Cleveland et al., 2006), although P addition did increase the cumulative CO_2_ emissions respired from the forest soil (Cleveland et al., 2002).

When we plotted RRs to experimental P addition using data from the studies from the literature review, the RRs of soil microbial respiration were consistently higher than the RRs of litter decomposition for the same amount of added P (Fig. 4). Thus, the contrasting response of soil microbial respiration and litter decomposition to P addition may be a common phenomenon. The effect of P addition on soil microbial respiration may have been previously overestimated. Instead, increased respiration rates may be partially attributed to the increase in abiotic C availability caused by P addition. The prevailing view that microbial activity in tropical forest soils is P-limited should be re-evaluated.

**Figure 4.**
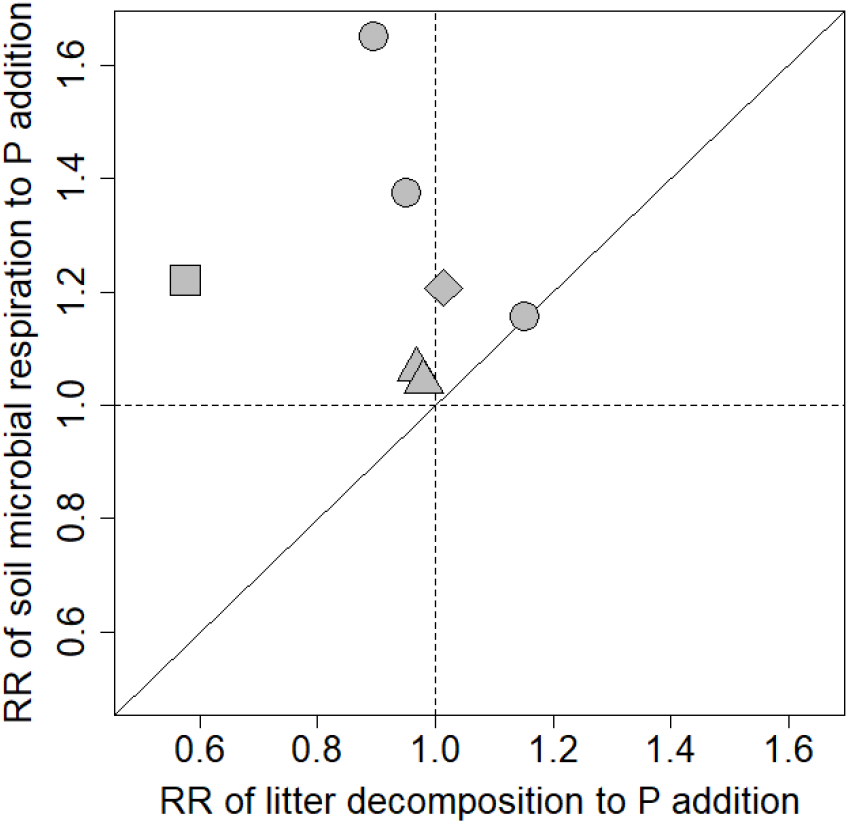
Response ratios (RRs) of soil microbial respiration and litter decomposition to experimental phosphorus (P) addition, calculated using data from previous studies. Filled circles represent data from studies of tropical montane forests in Hawaii (Hobbie and Vitousek, 2000; Reed et al., 2011). Filled diamonds represent data from studies of a tropical rain forest in Costa Rica (Cleveland et al., 2002; 2006). Filled squares represent data from a study of a tropical tree plantation in Indonesia (Mori et al., 2015). Filled triangles represent slightly revised data from a study of tropical tree plantations in Thailand (Mori et al., 2016).

## Acknowledgements

This study was financially supported by National Natural Science Foundation of China (41731176, 31670488, 4161101087), the Natural Science Foundation of Guangdong Province (2017A030313168), Grant-in-Aid for JSPS Postdoctoral Fellowships for Research Abroad (28, 601), and a grant from The Sumitomo Foundation (153082).

